# Inhibition of CPEB3 ribozyme elevates CPEB3 protein expression and polyadenylation of its target mRNAs, and enhances object location memory

**DOI:** 10.1101/2023.06.07.543953

**Authors:** Claire C. Chen, Joseph Han, Carlene A. Chinn, Jacob S. Rounds, Xiang Li, Mehran Nikan, Marie Myszka, Liqi Tong, Luiz F. M. Passalacqua, Timothy W. Bredy, Marcelo A. Wood, Andrej Lupták

## Abstract

A self-cleaving ribozyme that maps to an intron of the cytoplasmic polyadenylation element binding protein 3 (*CPEB3*) gene is thought to play a role in human episodic memory, but the underlying mechanisms mediating this effect are not known. We tested the activity of the murine sequence and found that the ribozyme’s self-scission half-life matches the time it takes an RNA polymerase to reach the immediate downstream exon, suggesting that the ribozyme-dependent intron cleavage is tuned to co-transcriptional splicing of the *CPEB3* mRNA. Our studies also reveal that the murine ribozyme modulates maturation of its harboring mRNA in both cultured cortical neurons and the hippocampus: inhibition of the ribozyme using an antisense oligonucleotide leads to increased CPEB3 protein expression, which enhances polyadenylation and translation of localized plasticity-related target mRNAs, and subsequently strengthens hippocampal-dependent long-term memory. These findings reveal a previously unknown role for self-cleaving ribozyme activity in regulating experience-induced co-transcriptional and local translational processes required for learning and memory.

**Significance Statement:** Cytoplasmic polyadenylation-induced translation is one of the key steps for regulating protein synthesis and neuroplasticity in the hippocampus. The CPEB3 ribozyme is a highly conserved mammalian self-cleaving catalytic RNA with unknown biological roles. In this study, we investigated how the intronic ribozyme affects the *CPEB3* mRNA maturation and translation, and its subsequent effect on memory formation. Our findings show that the ribozyme activity is anti-correlated with *CPEB3* mRNA splicing: inhibition of the ribozyme results in higher mRNA and protein levels, which contribute to long-term memory. Our studies offer new insights into the role of the CPEB3 ribozyme in neuronal translational control for the activity-dependent synaptic functions that underlie long-term memory and demonstrate a novel biological role for self-cleaving ribozymes.

## Introduction

Cytoplasmic polyadenylation element binding proteins (CPEBs) are RNA-binding proteins that modulate polyadenylation-induced mRNA translation, which is essential for the persistence of memory (Huang et al., 2003). CPEBs have been found in several invertebrate and vertebrate genomes, and four *CPEB* genes (*CPEB1–4*) have been identified in mammals (Si et al., 2003; Theis et al., 2003; Richter, 2007; Merkel et al., 2013; Afroz et al., 2014). All CPEB proteins have two RNA recognition domains (RRM motifs) and a ZZ-type zinc finger domain in the C-terminal region, but they differ in their N-terminal domains (Hake and Richter, 1994; Huang et al., 2006; Ivshina et al., 2014). *Aplysia* CPEB (ApCPEB), *Drosophila* Orb2, and mouse CPEB3 have two distinct functional conformations that correspond to soluble monomers and amyloidogenic oligomers, and have been implicated in the maintenance of long-term facilitation (LTF) in *Aplysia* and long-term memory in both *Drosophila* and mice (Miniaci et al., 2008; Si et al., 2010; Majumdar et al., 2012; Fioriti et al., 2015; Hervas et al., 2016; Rayman and Kandel, 2017; Hervas et al., 2020). In *Drosophila*, inhibition of amyloid-like oligomerization of Orb2 impairs the persistence of long-lasting memory, and deletion of the prion-like domain of Orb2 disrupts long-term courtship memory (Keleman et al., 2007; Hervas et al., 2016). The aggregated form of CPEB3, which is inhibited by SUMOylation, can mediate target mRNA translation at activated synapses (Drisaldi et al., 2015).

Following synaptic stimulation, CPEB3 interacts with the actin cytoskeleton, with a positive feedback loop of CPEB3/actin regulating remodeling of synaptic structure and connections (Stephan et al., 2015; Gu et al., 2020). Studies of CPEB3 in memory formation revealed that local protein synthesis and long-term memory storage are regulated by the prion-like CPEB3 aggregates, which are thought to strengthen synaptic plasticity in the hippocampus. While *CPEB3* conditional knockout mice display impairments in memory consolidation, object placement recognition, and long-term memory maintenance (Fioriti et al., 2015), global *CPEB3* knockout (*CPEB3*-KO) mice exhibit (i) enhanced spatial memory consolidation in the Morris water maze, (ii) elevated short-term fear memory in a contextual fear conditioning task, and (iii) improved long-term memory in a spatial memory task (water maze) (Chao et al., 2013). Moreover, dysregulation of translation of plasticity-associated proteins and post-traumatic stress disorder-like behavior after traumatic exposure is observed in *CPEB3*-KO mice (Lu et al., 2021).

In addition to encoding the CPEB3 protein, the mammalian *CPEB3* gene also encodes a functionally conserved self-cleaving ribozyme that maps to the second intron (Salehi-Ashtiani et al., 2006; Webb and Luptak, 2011; Bendixsen et al., 2021) (Fig. 1A). Several mammalian ribozymes have been identified (Salehi-Ashtiani et al., 2006; Martick et al., 2008; de la Pena and Garcia-Robles, 2010; Perreault et al., 2011; Hernandez et al., 2020; Chen et al., 2021), including the highly active sequence in the *CPEB3* gene. The CPEB3 ribozyme belongs to hepatitis delta virus (HDV)-like ribozymes, which are self-cleaving RNAs widespread among genomes of eukaryotes, bacteria, and viruses (Webb et al., 2009; Eickbush and Eickbush, 2010; Ruminski et al., 2011; Sanchez-Luque et al., 2011; Weinberg et al., 2015; Passalacqua et al., 2017). The biological roles of these ribozymes vary widely and include processing rolling-circle transcripts during HDV replication (Sharmeen et al., 1988; Wu et al., 1989), 5′-cleavage of retrotransposons (Eickbush and Eickbush, 2010; Ruminski et al., 2011; Sanchez-Luque et al., 2011), and in one bacterial example, the HDV-like ribozyme may mediate metabolite-dependent regulation of gene expression (Passalacqua et al., 2017). Furthermore, the genomic locations of these catalytic RNAs suggest that they are involved in many other biological processes. Recent analysis suggests that, although the biological function remains unknown, CPEB3 ribozymes have had a role in mammals for over 100 million years (Bendixsen et al., 2021). In humans, a single nucleotide polymorphism (SNP) at the ribozyme cleavage site leads to a 3-fold higher rate of *in vitro* self-scission, which correlates with poorer performance in an episodic memory task (Salehi-Ashtiani et al., 2006; Vogler et al., 2009) and suggests that the ribozyme activity may play a role in memory formation.

**Figure 1.**
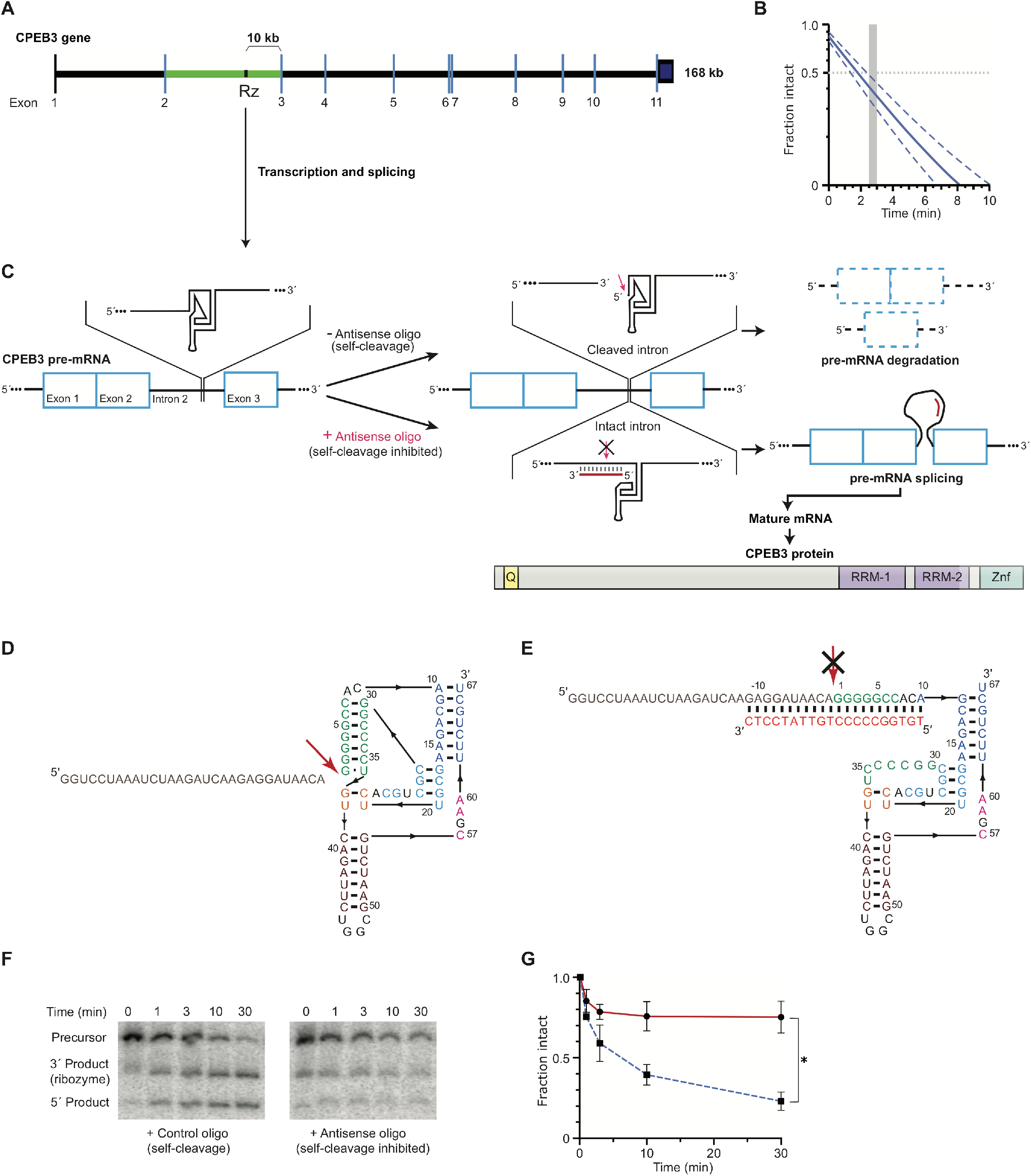
*CPEB3* gene structure and activity of its intronic self-cleaving ribozyme. ***A***, Schematic representation of mouse *CPEB3* gene. Rz denotes the location of the self-cleaving ribozyme in the 2^nd^ intron (green) between the 2^nd^ and 3^rd^ exons. ***B***, Co-transcriptional self-cleavage activity of a 470-nt construct, incorporating the 72-nt ribozyme, which cuts the transcript 233 nts from the 5′ terminus (see Table 1 for kinetic parameters of this and other constructs). Log-linear graph of self-cleavage is shown with solid blue line (dashed lines show ± standard deviation). Gray dotted line indicates mid-point of self-cleavage (with resulting *t*_1/2_ of ∼2 min). Gray bar indicates the approximate time range for RNAPII to travel from the ribozyme to the 3^rd^ exon, at which point ∼40% of the intron would remain intact. ***C***, Inhibition of the CPEB3 ribozyme by an ASO targeting its cleavage site and the resulting effect on the levels of the spliced mRNA and the encoded protein. ***D***, Secondary structure of the ribozyme (colored by structural elements (Webb and Luptak, 2011)). Sequence upstream of the ribozyme is shown in gray, and the site of self-scission is shown with a red arrow. ***E***, Model of the ribozyme inhibited by the ASO (red letters) showing base-pairing between the ASO and 10 nts upstream and downstream of the ribozyme cleavage site. Inhibition of self-scission is indicated by crossed arrow (***C*** and ***E***). ***F***, Inhibition of CPEB3 ribozyme self-scission *in vitro* in the presence of ASO. Scrambled or antisense oligonucleotides (1 µM) were added during co-transcriptional self-cleavage reactions. ***G***, Fraction intact values were calculated and plotted vs. time. Significant inhibition of co-transcriptional self-scission by the ASO (red line, compared with control oligo shown in blue), resulting in increase of intact RNA (***F*** and ***G***), is observed at the 3-min time point relevant to the transcription of the *CPEB3* gene (***A*** and ***B***).

**Table 1.** Kinetic parameters of murine CPEB3 ribozyme constructs^2^

| Construct<sup>1</sup> | A | $k_1$ | B | $k_2$ | C |
| --- | --- | --- | --- | --- | --- |
| -10/72 | $0.72 \pm 0.09$ | $0.39 \pm 0.09$ | | | $0.082 \pm 0.026$ |
| -49/72/165 | $0.88 \pm 0.02$ | $0.42 \pm 0.04$ | $0.013 \pm 0.015$ | $0.11 \pm 0.03$ | $0.04 \pm 0.02$ |
| -233/72/165 | $0.78 \pm 0.04$ | $0.31 \pm 0.04$ | $0.035 \pm 0.006$ | $0.17 \pm 0.02$ | $0.029 \pm 0.005$ |

<sup>1</sup> Construct size is defined as (length of sequence upstream of the ribozyme cleavage site)/[CPEB3 ribozyme (72 nts)]/(downstream sequence).
<sup>2</sup> Co-transcriptional self-scission was modeled by a bi-exponential decay model with a residual. A and B represent fractions of the population cleaving with fast ( $k_1$ ) and slow ( $k_2$ ) rate constants, respectively. The residual (C) is interpreted as a fraction of the population that does not self-cleave. Errors represent SEM of at least three experiments. For the smallest ribozyme construct (-10/72), a monoexponential decay function was sufficient to model the data.

While the CPEB3 protein is well established as a modulator of memory formation and learning, the molecular and physiological functions of the intronic CPEB3 ribozyme have not been tested. Using synthetic ribozymes placed within introns of mammalian genes, previous work showed that splicing of the surrounding exons is sensitive to the continuity of the intron: fast ribozymes caused efficient self-scission of the intron, leading to unspliced mRNA and lower protein expression. In contrast, slow ribozymes had no effect on mRNA splicing and subsequent protein expression (Fong et al., 2009). Based on this observation, we tested the hypothesis that inhibition of the CPEB3 ribozyme co-transcriptional self-scission will promote *CPEB3* mRNA splicing (Fig. 1A) and increase the expression of full-length mRNA and CPEB3 protein, leading to polyadenylation of its target mRNAs and enhancement in the consolidation of hippocampal-dependent memory.

## Materials and Methods

### Primary cortical neuronal culture

Pregnant female C57BL/6 mice (The Jackson Laboratory) were euthanized at E18, and embryos were collected into an ice-cold Neurobasal medium (Thermo Fisher Scientific). Embryonic cortices were dissected, meninges were removed, and tissues were minced. Cells were mechanically dissociated, passed through a 40-µm cell strainer, counted, and plated at a density of 0.5 × 10^6^ cells per well in six-well plates coated with poly-D-lysine (Sigma-Aldrich). Neuronal cultures were maintained at 37 °C with 5% CO_2_, and grown in Neurobasal medium containing 2% B27 supplement (Thermo Fisher Scientific), 1% penicillin/streptomycin (Thermo Fisher Scientific), and 2 mM L-glutamine (Thermo Fisher Scientific) for 7–10 days *in vitro* (DIV), with 50% of the medium being replaced every 3 days. All experimental procedures were performed according to the National Institutes of Health Guide for the Care and Use of Laboratory Animals and approved by the Institutional Animal Care and Use Committee of the University of California, Irvine.

### Mice

C57BL/6J mice (8–10 weeks old, The Jackson Laboratory) were housed in a 12-h light/dark cycle and had free access to water and food. All experiments were conducted during the light cycle. All experimental procedures were performed according to the National Institutes of Health Guide for the Care and Use of Laboratory Animals and approved by the Institutional Animal Care and Use Committee of the University of California, Irvine.

### Measurement of co-transcriptional self-scission of the CPEB3 ribozyme

*In vitro* co-transcriptional cleavage kinetics were measured using a previously described method that utilizes standard T7 RNA polymerase *in vitro* transcription under minimal MgCl_2_ concentration, followed by a 25-fold dilution of the reaction to stop the synthesis of transcripts and to allow the study of the self-scission reaction without the need for purification or additional preparation steps (Passalacqua et al., 2017). Transcription reactions were set up in a 5 µL volume and incubated for 10 minutes at 24 °C. The reactions contained the following components: 1 µL of 5× transcription buffer (10 mM spermidine, 50 mM dithiothreitol, 120 mM Tris chloride buffer, pH 7.5, and 0.05% Triton X-100), 1 µL of 5× ribonucleoside triphosphates (final total concentration of 6.8 mM), 1 µL of 5 mM Mg^2+^, 1 µL DNA amplified by PCR to about 1 µM final concentration, 0.5 µL of 100% DMSO, 0.15 µL of water, 0.1 µL of murine RNase inhibitor (40,000 units/mL, New England Biolabs), 0.125 µL of T7 polymerase, and 0.125 µL [α-^32^P]ATP. To prevent initiation of new transcription, the reactions were diluted into 100 µL of physiological-like buffer solution at 37 °C. The solution consisted of 2 mM Mg^2+^ (to promote ribozyme self-scission), 140 mM KCl, 10 mM NaCl, and 50 mM Tris chloride buffer (pH 7.5). The 100 µL solution was then held at 37 °C for the reminder of the experiment while aliquots were withdrawn at various time points. An equal volume of 4 mM EDTA/7 M urea stopping solution was added to each aliquot collected. Aliquots were resolved using denaturing polyacrylamide gel electrophoresis (PAGE, 7.5% polyacrylamide, 7 M urea). The PAGE gel was exposed to a phosphorimage screen for ∼2 hours and analyzed using a Typhoon imaging system (GE Healthcare). Band intensities corresponding to the uncleaved ribozymes and the two products of self-scission were analyzed using ImageQuant (GE Healthcare) and exported into Excel. Fraction intact was calculated as the intensity of the band corresponding to the uncleaved ribozyme divided by the sum of band intensities in a given PAGE lane. The data were fit to a biexponential decay model:

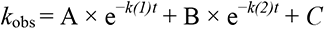

In the case of the smallest (minimal) murine CPEB3 ribozyme construct (-10/72; Table 1), the data were modeled by a monoexponential decay with an uncleaved fraction (using parameters A, *k*_1_, and C only).

### *In vitro* co-transcriptional cleavage kinetics in the presence of antisense oligonucleotides (ASO)

To test inhibition of the CPEB3 ribozyme by antisense oligos (ASOs), *in vitro* transcription was performed in a solution containing 10 mM dithiothreitol (DTT), 2 mM spermidine, 4.5 mM MgCl_2_; GTP, UTP, and CTP (1.25 mM each); 250 µM ATP; 4.5 µCi of [α-^32^P]ATP (PerkinElmer); 40 mM HEPES (pH 7.4), and 1 unit of T7 RNA polymerase. A 5.0 µL transcription reaction was initiated by the addition of 0.5 pmol of DNA template, and the mixture was incubated at 24 °C for 10 min. A 1.0 µL aliquot of the reaction was withdrawn, and its transcription and self-scission were terminated by the addition of urea loading buffer. The remaining 4.0 µL volume was diluted 25-fold (final volume of 100 µL) into a physiological-like solution [50 mM HEPES buffer (pH 7.4), 10 mM NaCl, 140 mM KCl, 10 mM MgCl_2_, and 1 µM of the ASO of interest] at 37 °C. A control experiment was performed in the presence of scrambled ASO. 5 µL aliquots were collected at the indicated times and terminated by the addition of 5 µL denaturing loading buffer (20 mM EDTA, 8 M urea, and the loading dyes xylene cyanol and bromophenol blue). Samples were resolved on a 10% polyacrylamide gel electrophoresis (PAGE) under denaturing conditions (7 M urea). The PAGE gel was exposed to a phosphorimage screen and analyzed using Typhoon phosphorimager and ImageQuant software (GE Healthcare). Band intensities were analyzed by creating line profiles of each lane using ImageQuant. Self-cleavage data were fit to a monoexponential decay function:

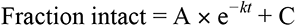

Where A represents the relative fractions of the ribozyme population cleaving with an apparent rate constant *k*, and C represents the population remaining uncleaved. The model was fit to the data using a linear least-squares analysis and the Solver module of Microsoft Excel.

### Antisense oligonucleotides (ASOs)

ASOs used in this study are 20 nucleotides in length and are chemically modified with 2′-*O*-methoxyethyl (MOE, underlined) and 2′,4′-constrained ethyl (cEt, bold) (Seth et al., 2009). All internucleoside linkages are modified with phosphorothioate linkages to improve nuclease resistance. ASOs were solubilized in sterile phosphate-buffered saline (PBS). The sequences of the ASOs are as follows (all cytosine nucleobases are 5-methyl-substituted):

Scrambled control ASO: 5′-**C**<u>CT</u>**T**<u>CC</u>**C**<u>TG</u>**A**<u>AG</u>**G**<u>TT</u>C<u>CT</u>**C**<u>C</u>-3′;

CPEB3 ribozyme ASO: 5′-**T**<u>GT</u>**G**<u>GC</u>**C**<u>CC</u>**C**<u>TG</u>**T**<u>TA</u>**T**<u>CC</u>**T**<u>C</u>-3′.

### Neuronal stimulation

Neurons were treated with ASO or scrambled ASO (1 µM) for 18 hours prior to neuronal stimulation. To study activity-dependent gene regulation, neuronal cultures were treated with vehicle, 5 µM glutamate (10 minutes), or 35 mM KCl (5 minutes). After stimulation, cultures were washed with Hanks’ buffered salt solution (HBSS, Thermo Fisher Scientific), and then fresh medium was added.

### Quantitative RT-PCR analysis

Total RNA was isolated from primary cortical neurons or mouse hippocampus using TRI reagent (Sigma-Aldrich) according to the manufacturer’s protocol. RNA concentration was measured using a NanoDrop ND-1000 spectrophotometer (Thermo Fisher Scientific). Total RNA was reverse transcribed using random decamers and M-MLV reverse transcriptase (Promega)/Superscript II RNase H reverse transcriptase (Thermo Fisher Scientific). Quantitative RT-PCR was performed on a BioRad CFX Connect system using iTaq Universal SYBR Green Supermix (BioRad). Designed primers were acquired from Integrated DNA Technologies and are provided in Table 2. Desired amplicons were verified by melting curve analysis and followed by gel electrophoresis. The starting quantity of DNA from each sample was determined by interpolation of the threshold cycle (CT) from a standard curve of each primer set. Relative gene expression levels were normalized to the endogenous gene *GAPDH*.

**Table 2.**
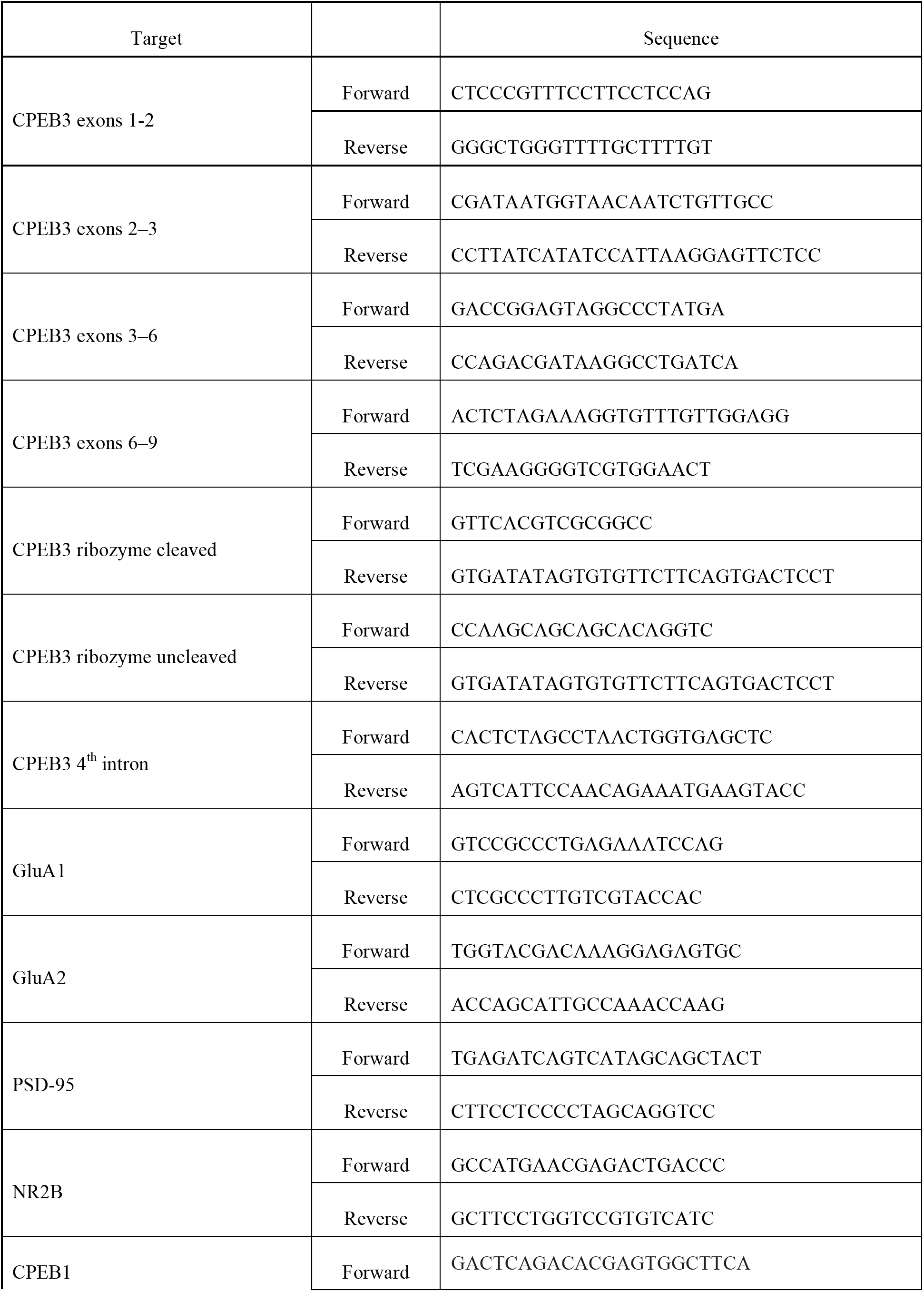
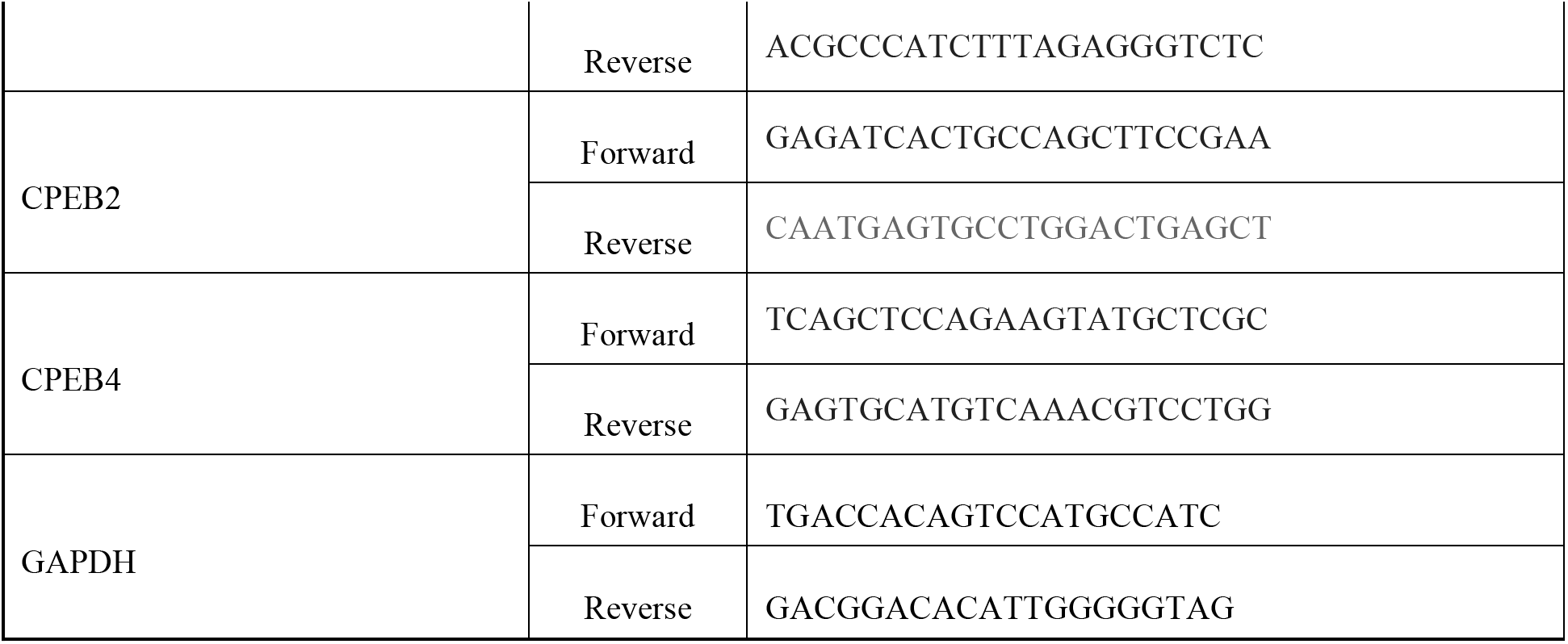
Primers used in qPCR

### Immunoblotting

Primary cortical neurons or mouse hippocampal tissues were lysed in RIPA lysis buffer with protease inhibitor (Santa Cruz Biotechnology). Crude synaptosomal fractions were prepared as previously described (Wirths, 2017). Protein concentrations were measured using bicinchoninic acid (BCA) protein assay (Thermo Fisher Scientific). Protein samples (10–30 µg) were loaded on 10% sodium dodecyl sulfate polyacrylamide (SDS-PAGE) gels and separated by electrophoresis. Gels were electro-transferred onto polyvinylidene fluoride (PVDF) membranes using a semi-dry transfer system (BioRad). Membranes were either blocked with 5% nonfat milk or 5% bovine serum albumin (BSA) in Tris-buffered saline/Tween 20 (0.1% [vol/vol]) (TBST) for 1 hour at room temperature. Membranes were incubated with primary antibodies overnight at 4 °C. After primary antibody incubation, membranes were washed three times with TBST and then incubated with secondary antibodies for 1 hour at room temperature. Bands were detected using an enhanced chemiluminescence (ECL) kit (Thermo Fisher Scientific), visualized using BioRad Chemidoc MP imaging system, and analyzed by Image Lab software (BioRad). GAPDH was used as a loading control.

The following antibodies were used: anti-CPEB3 (Abcam, 1:1000); anti-GluA1 (UC Davis/NIH NeuroMab Facility, 1:1000); anti-GluA2 (Proteintech, 1:2000); anti-PSD95 (Proteintech, 1:2000); anti-NR2B (Proteintech, 1:2000); anti-CPEB1 (ABclonal, 1:1000), CPEB4 (Proteintech, 1:1000); anti-GAPDH (Proteintech, 1:10,000); donkey anti-rabbit-HRP (Thermo Fisher Scientific, 1:10,000); goat anti-mouse-HRP (R&D system, 1:1000).

### *In vitro* XTT cell viability assay

Primary cortical neurons (10,000 to 20,000 cells/well) were plated onto 96-well plates coated with poly-D-lysine. After 7–14 days, ASOs or scrambled ASOs were added, and the resulting solutions were incubated for 18 hours. Cell viability was determined using the 2,3-bis[2-methoxy-4-nitro-5-sulfophenyl]-2H-tetrazolium-5-carboxyanilide inner salt (XTT) assay according to the manufacturer’s protocol (Biotium). The assay utilizes the ability of viable cells with active metabolism to reduce the yellow tetrazolium salt to the soluble orange formazan product using mitochondrial dehydrogenase enzymes. The XTT reagent was added to each well and incubated for 2–4 hours at 37 °C and under 5% CO_2_. Absorbance was measured at 450 nm with a reference wavelength of 680 nm using a Biotek Synergy HT microplate reader. Results were normalized to control, and all samples were assayed in triplicate.

### Stereotaxic surgeries

C57/BL6J mice (8–10 weeks old, Jackson Laboratory), housed under standard conditions with light-control (12 hour light/12 hour dark cycles), were anaesthetized with an isoflurane (1–3%)/oxygen vapor mixture. Mice were infused bilaterally to the CA1 region of the dorsal hippocampus with ribozyme ASO or scrambled ASO diluted in sterile PBS. The following coordinates were used, relative to bregma: medial-lateral (ML), ±1.5 mm; anterior-posterior (AP), −2.0 mm; dorsal-ventral (DV), −1.5 mm. ASOs or vehicle (1 nmol/µL) were infused bilaterally at a rate of 0.1 µL/min using a Neuros Hamilton syringe (Hamilton company) with a syringe pump (Harvard Apparatus). The injectors were left in place for 2 minutes to allow diffusion, and then were slowly removed at a rate of 0.1 mm per 15 sec. The incision site was sutured, and mice were allowed to recover on a warming pad and then were returned to cages. For all surgeries, mice were randomly assigned to the different conditions to avoid grouping same treatment conditions in time.

### Object location memory (OLM) tasks

The OLM task was performed to assess hippocampus-dependent memory, as previously described (Vogel-Ciernia and Wood, 2014). Briefly, naïve C57/BL6J mice (8–12 weeks old; n = 10–12/group; CPEB3 ribozyme ASO or scrambled ASO) were trained and tested. Prior to training, mice were handled 1–2 minutes for 5 days and then habituated to the experimental apparatus for 5 minutes on 6 consecutive days in the absence of objects. During training, mice were placed into the apparatus with two identical objects and allowed to explore the objects for 10 minutes. Twenty-four hours after training, mice were exposed to the same arena, and long-term memory was tested for 5 minutes, with the two identical objects present, one of which was placed in a novel location. For all experiments, objects and locations were counterbalanced across all groups to reduce bias. Videos of training and testing sessions were analyzed for discrimination index (DI) and total exploration time of objects. The videos were scored by observers blind to the treatment. The exploration of the objects was scored when the mouse’s snout was oriented toward the object within a distance of 1 cm or when the nose was touching the object. The relative exploration time was calculated as a discrimination index (DI = (*t*_novel_ – *t*_familiar_) / (*t*_novel_ + *t*_familiar_) × 100%). Mice that demonstrated a location or an object preference during the training trial (DI > ±20) were removed from analysis.

### 3’ RACE

Total RNA was extracted from the mouse CA1 hippocampus, and 3*’* rapid amplification of cDNA ends (3*’* RACE) was performed to study the alternative polyadenylation. cDNA was synthesized using oligo(dT) primers with 3*’* RACE adapter primer sequence at the 5*’* ends. This cDNA library results in a universal sequence at the 3*’* end. A gene-specific primer (GSP) and an anchor primer that targets the poly(A) tail region were employed for the first PCR using the following protocol: 95 °C for 3 minutes, then 30 cycles of 95 °C for 30 seconds, 55 °C for 30 seconds, and 72 °C for 3 minutes, with a final extension of 72 °C for 5 minutes. To improve specificity, a nested PCR was then carried out using primers internal to the first two primers. Upon amplification condition optimization, a quantitative PCR was performed on the first diluted PCR product using the nested primers, and a standard curve of the primer set was generated to measure the relative expression of 3*’*-mRNA and alternative polyadenylation. All primers used in this study are listed in Table 3. When resolved using agarose gel electrophoresis, this nested-primer qPCR produced single bands corresponding to the correct amplicons of individual cDNAs.

**Table 3.**
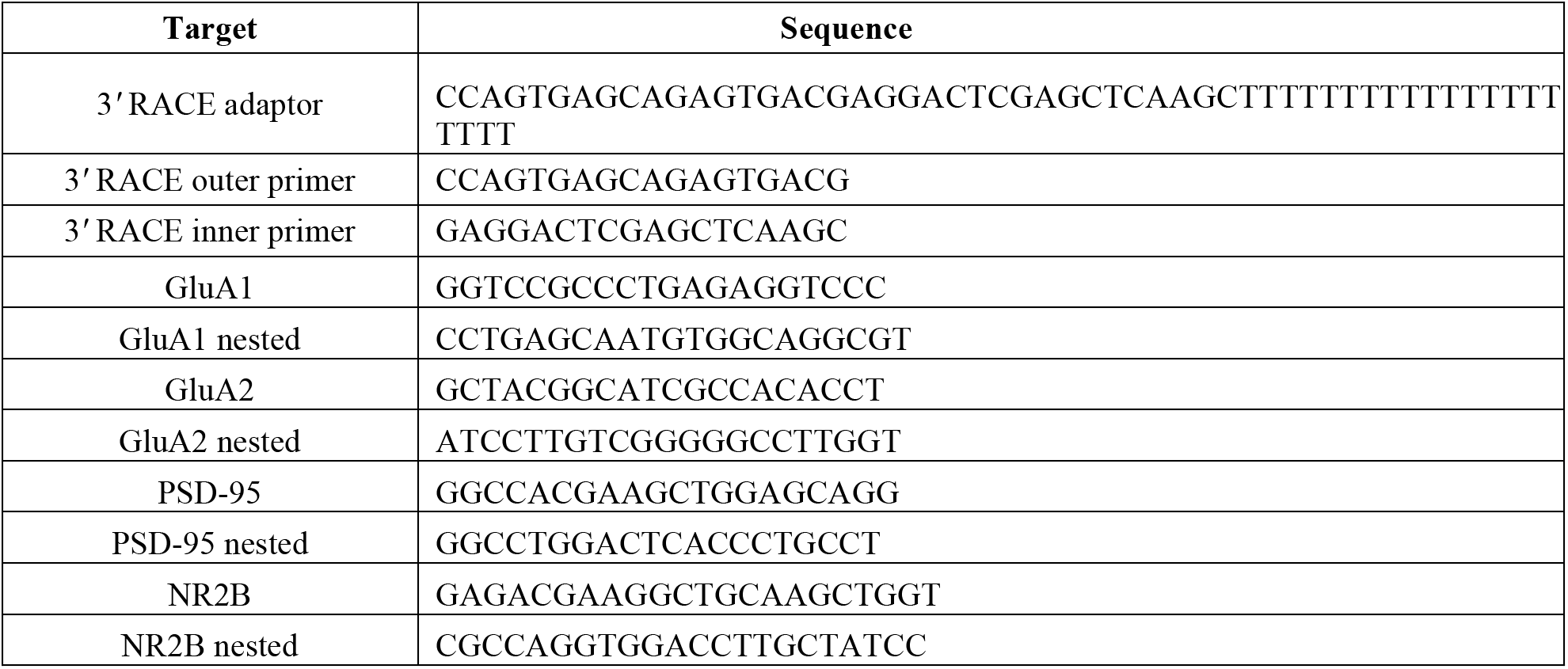
Primers used in 3*’* RACE

### Statistical analysis

Data are presented as means ± SEM. Statistical analyses were performed using GraphPad Prism (GraphPad Prism Software). Statistical differences were determined using (i) two-tailed Welch’s *t* test when comparing between 2 independent groups, (ii) one-way ANOVA with Sidak’s *post hoc* tests when comparing across 3 or more independent groups, and (iii) two-way ANOVA with Sidak’s *post hoc* tests when comparing two factors. *P* < 0.05 was considered significant.

## Results

### Antisense oligonucleotides (ASOs) inhibit CPEB3 ribozyme self-scission

To determine whether the CPEB3 ribozyme activity modulates expression of the CPEB3 protein by disrupting co-transcriptional splicing of the *CPEB3* mRNA, we started by measuring the co-transcriptional self-scission of the murine variant of the ribozyme *in vitro* and determined the half-life (*t*_1/2_) to be ∼2–3 minutes (Fig. 1B and Table 1). This rate of self-scission is similar to that measured previously for chimp and fast-reacting human variants of the ribozyme (Chadalavada et al., 2010). Because the distance from the ribozyme cleavage site to the 3^rd^ exon in the *CPEB3* gene is 9931 nucleotides (Fig. 1A) and the RNA polymerase II (RNAPII) transcription rate of long mammalian genes is estimated to be ∼3.5–4.1 knt/min (Singh and Padgett, 2009), RNAPII should require about 2.5–3 minutes to travel from the ribozyme to the 3^rd^ exon. The nascent ribozyme thus self-cleaves in about the same time as it takes the RNAPII to synthesize the remaining part of the intron and the next exon, at which point the splicing machinery is expected to mark the intron–exon junction. This observation suggests that the ribozyme activity is tuned to the co-transcriptional processing of the *CPEB3* pre-mRNA: a significantly faster rate of self-scission would lead to a high fraction of cleaved, unspliced pre-mRNAs, whereas slow self-cleavage rate would have no effect on the *CPEB3* pre-mRNA splicing.

ASOs are synthetic single-stranded nucleic acids that can bind to pre-mRNA or mature RNA by base-pairing, and typically trigger RNA degradation by RNase H. ASOs have also been employed to modulate alternative splicing, suggesting that they act co-transcriptionally *in vivo* [e.g., to correct the *SMN2* gene (Hua et al., 2010)]. We designed and screened a series of ASOs with the goal of blocking co-transcriptional self-scission of the CPEB3 ribozyme. The greatest inhibition was observed when the ASO was bound to the ribozyme cleavage site (Fig. 1C, D, E); similar ASOs have been used to inhibit *in vitro* co-transcriptional self-scission of other HDV-like ribozymes (Harris et al., 2004; Webb et al., 2009). As the CPEB3 ribozyme was synthesized, 80% of it remained uncleaved in the presence of this ASO, compared to 20% in the presence of a control oligonucleotide at the 30-min time point (unpaired *t* test, *t*_(3.599)_ = 8.204, *P* = 0.0019; Fig.1F, G). This ASO and a scrambled control sequence were used in all subsequent *in cellulo* and *in vivo* experiments.

### *CPEB3* mRNA expression and ribozyme activity are upregulated in response to neuronal stimulation

Neuronal activity-dependent gene regulation is essential for synaptic plasticity (Neves et al., 2008). To investigate the effect of the CPEB3 ribozyme on *CPEB3* mRNA expression and measure its effect on maturation and protein levels, we began by stimulating primary cortical neurons with glutamate or potassium chloride (KCl). *CPEB3* mRNA levels were measured using primers that specifically amplified exon–exon splice junctions (Exons 2–3, 3– 6, 6–9; Fig. 1A). We found that membrane depolarization by KCl led to an upregulation of *CPEB3* mRNA 2 hours post-stimulation, compared with non-stimulated cultures (exons 2–3: *F*_(5,12)_ = 18.02, *P* < 0.0001; exons 3–6: *F*_(5,12)_ = 25.48, *P* < 0.0001; exons 6–9: *F*_(5,12)_ = 4.376, *P* = 0.0168; one-way ANOVA with Sidak’s *post hoc* tests; Fig. 2A). To examine CPEB3 ribozyme activity, total ribozyme and uncleaved ribozyme levels were measured by qRT-PCR, which showed that ribozyme expression is elevated at 1 hour following KCl treatment (*F*_(5,17)_ = 12.96, *P* < 0.0001; one-way ANOVA with Sidak’s *post hoc* tests; Fig. 2B). Similarly, glutamate stimulation resulted in increased expression of spliced exons by 2–3 fold at 2 hours, with a decrease observed at later time points (exons 2–3: *F*_(5,21)_ = 5.826, *P* = 0.0016; exons 3–6: *F*_(5,22)_ = 2.002, *P* = 0.1181; exons 6–9: *F*_(5,22)_ = 1.763, *P* = 0.1622; one-way ANOVA with Sidak’s *post hoc* tests; Fig. 2C), and increased ribozyme expression correlated with *CPEB3* mRNA expression (*F*_(5,26)_ = 4.657, *P* = 0.0036; one-way ANOVA with Sidak’s *post hoc* tests; Fig. 2D). This finding is supported by previous studies showing that synaptic stimulation by glutamate leads to an increase in CPEB3 protein expression in hippocampal neurons (Fioriti et al., 2015) and that treatment with kainate likewise induces CPEB3 expression in the hippocampus (Theis et al., 2003). The cleaved fraction of the ribozyme was greatest at the highest point of *CPEB3* mRNA expression, indicating efficient co-transcriptional self-scission. Together, these data indicate that the self-cleaving CPEB3 ribozyme is expressed—and potentially activated—in response to neuronal activity, and suggest that CPEB3 ribozyme *cis*-regulates the maturation of CPEB3 mRNA.

**Figure 2.**
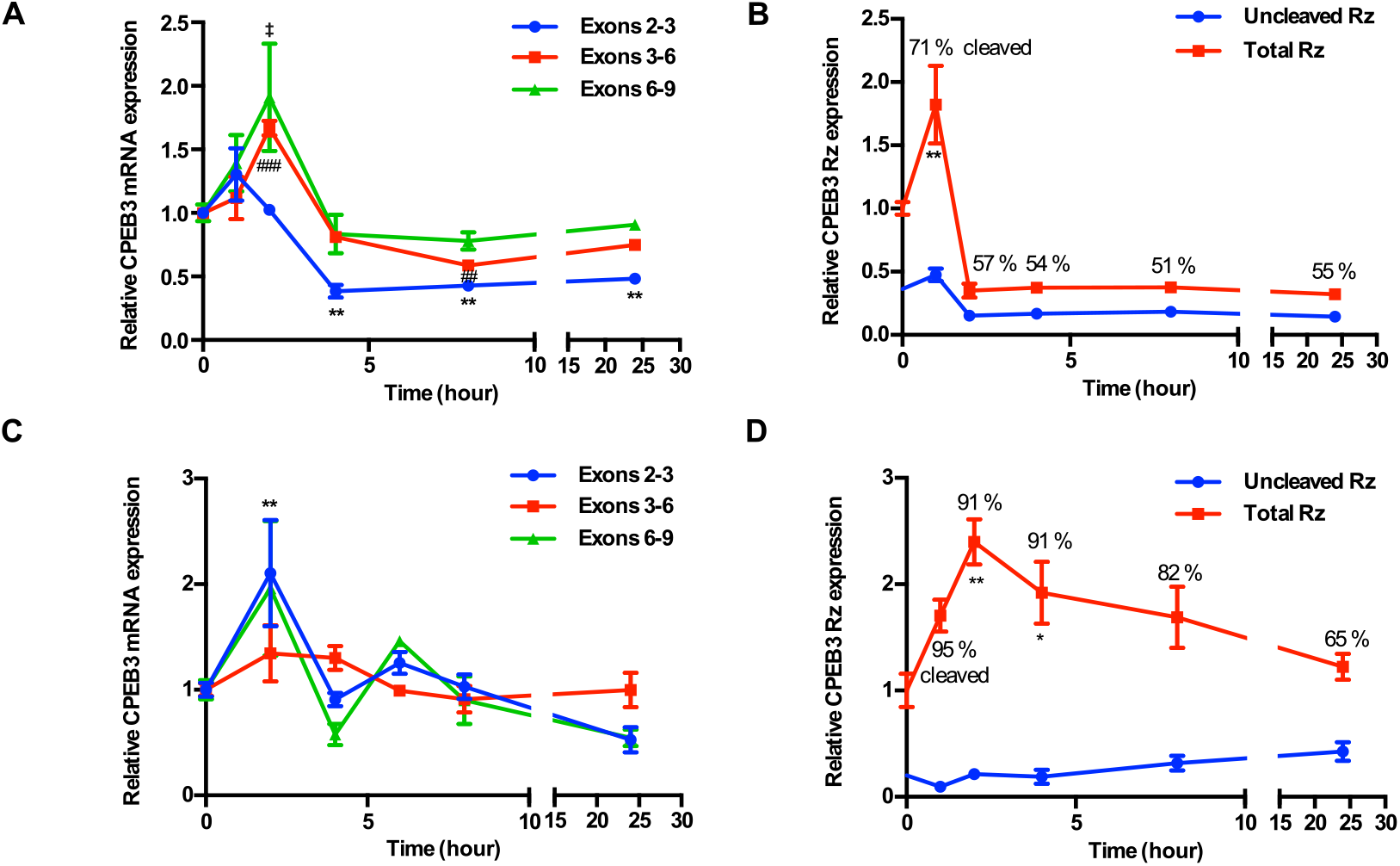
*CPEB3* expression in primary cortical neurons (DIV14). ***A***, KCl stimulation profile of the *CPEB3* gene showing induction of spliced CPEB3 exons. ***B***, KCl stimulation profile of CPEB3 ribozyme expression (uncleaved and total). Cleaved ribozyme fraction is calculated as [(total ribozyme – uncleaved ribozyme)/total ribozyme] and shown as % cleaved. ***C***, Expression of *CPEB3* mRNA exons 2–3 is upregulated 2 hours after glutamate stimulation. ***D***, Glutamate stimulation induces an increase in CPEB3 ribozyme levels at 2-hour time point. \**P* < 0.05, \*\**P* < 0.01, ##*P* < 0.01, ###*P* < 0.001, ‡*P* < 0.05. Data are presented as mean ± SEM.

### *CPEB3* mRNA levels increase in primary neuronal cultures treated with ribozyme inhibitor

Because our data showed that CPEB3 ribozyme expression is correlated with mRNA expression, we hypothesized that modulation of the ribozyme activity may alter *CPEB3* mRNA splicing. If so, then abrogation of the ribozyme self-scission would result in uncleaved second intron and higher levels of spliced mRNA. We inhibited the ribozyme using ASOs that were designed to increase thermal stability of complementary hybridization and, as a result, to induce higher binding affinity for the ribozyme. To study the effect of the CPEB3 ribozyme on *CPEB3* mRNA expression, neuronal cultures were pretreated with either an ASO or a non-targeting (scrambled) control oligonucleotide, followed by KCl stimulation. In the absence of ASO, KCl induced a rapid and robust increase in ribozyme levels compared to cultures containing scrambled ASO, and this effect was suppressed in the presence of ASO, which is consistent with the ASO blocking the ribozyme (two-way ANOVA with Sidak’s *post hoc* tests, significant main effect of KCl: *F*_(1,19)_ = 8.058, *P* = 0.0105; significant effect of ASO: *F*_(1,19)_ = 12.88, *P* = 0.0020; no significant interaction: *F*_(1,19)_ = 3.557, *P* = 0.0747; Fig. 3A). At an early time point (2 hours post-KCl induction), the ASO-containing culture displayed an increase of spliced mRNA (Exons 2–3: two-way ANOVA with Sidak’s *post hoc* tests, significant effect of ASO: *F*_(1,20)_ = 21.81, *P* = 0.0001, no significant effect of KCl: *F*_(1,20)_ = 0.1759, *P* = 0.6794; no significant interaction: *F*_(1,20)_ = 0.001352, *P* = 0.9710; Fig. 3B. Exons 3–6: two-way ANOVA with Sidak’s *post hoc* tests, significant ASO × KCl interaction: *F*_(1,19)_ = 5.726, *P* = 0.0272; significant effect of ASO: *F*_(1,19)_ = 8.042, *P* = 0.0106; no significant effect of KCl: *F*_(1,19)_ = 0.2922, *P* = 0.5951; Fig. 3C. Exons 6–9: two-way ANOVA with Sidak’s *post hoc* tests, no significant effect of KCl: *F*_(1,19)_ = 1.218, *P* = 0.2835, no significant effect of ASO: *F*_(1,19)_ = 3.919, *P* = 0.0624, and no significant interaction: *F*_(1,19)_ = 0.002317, *P* = 0.9621; Fig. 3D). The ASO likely prevents CPEB3 ribozyme from cleaving the intron co-transcriptionally and thereby promotes mRNA maturation. At 24 hours post-KCl induction, we observed no significant difference in CPEB3 ribozyme expression among groups (two-way ANOVA with Sidak’s *post hoc* tests, no significant effect of KCl: *F*_(1,18)_ = 0.7897, *P* = 0.3859, no significant effect of ASO: *F*_(1,18)_ = 0.03687, *P* = 0.8499, and no significant interaction: *F*_(1,18)_ = 0.9533, *P* = 0.3418; Fig. 3E). Likewise, the level of *CPEB3* mRNA exons 2–3 returned to the basal level (two-way ANOVA with Sidak’s *post hoc* tests, no significant effect of KCl: *F*_(1,19)_ = 0.0004856, *P* = 0.9826; no significant effect of ASO: *F*_(1,19)_ = 3.188, *P* = 0.0902, and no significant interaction: *F*_(1,19)_ = 0.4343, *P* = 0.5178; Fig. 3F), while exons 3– 6 remained slightly elevated in the ASO-treatment groups (two-way ANOVA with Sidak’s *post hoc* tests, significant effect of ASO: *F*_(1,19)_ = 11.48, *P* = 0.0031; no significant effect of KCl: *F*_(1,19)_ = 2.252, *P* = 0.1499; no significant interaction: *F*_(1,19)_ = 0.04047, *P* = 0.8417; Fig. 3G). The mRNA expression of CPEB3 exons 6–9 remained stable over time and was not affected by ASO treatment or KCl stimulation (two-way ANOVA with Sidak’s *post hoc* tests, no significant effect of KCl: *F*_(1,19)_ = 0.6316, *P* = 0.4366; no significant effect of ASO: *F*_(1,19)_ = 1.364, *P* = 0.2573, and no significant interaction: *F*_(1,19)_ = 0.1475, *P* = 0.7052; Fig. 3H). We further evaluated whether inhibition of CPEB3 ribozyme affects the levels of full-length *CPEB3* mRNA, and we found that ASO treatment led to a significant increase of spliced exons 2–9 (which correspond to the protein-coding segment of the mRNA) at the 2-hour time point (unpaired *t* test, *t*_(10.00)_ = 3.774, *P* = 0.0036; Fig. 3I). Taken together, these data show that the CPEB3 ribozyme modulates the production of the full-length *CPEB3* mRNA.

**Figure 3.**
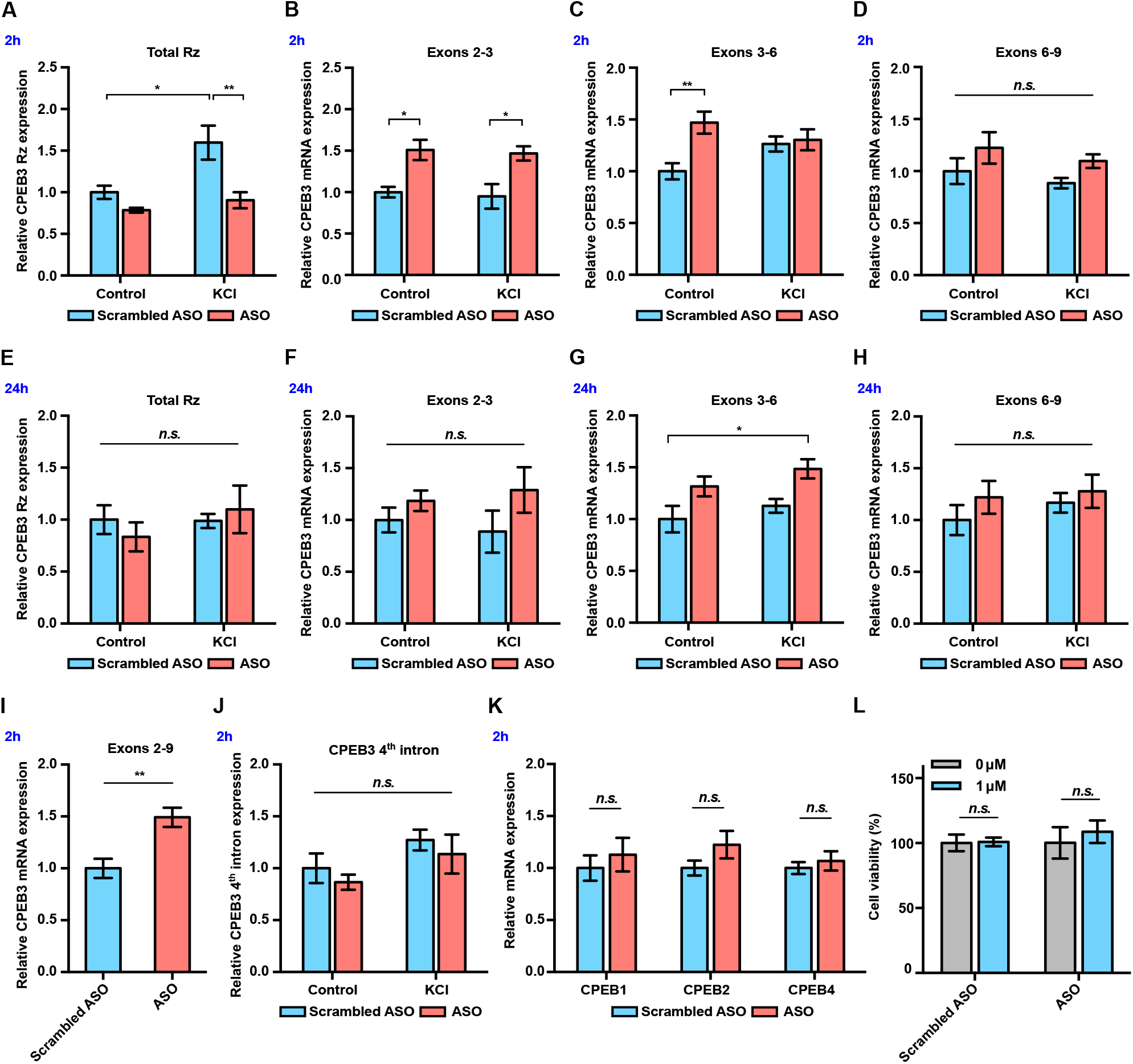
*CPEB3* mRNA is upregulated in primary neuronal cultures (DIV14) treated with ribozyme ASO. ***A***, CPEB3 ribozyme levels increase together with levels of the surrounding exons 2 hours post-stimulation in experiments with control ASO. Ribozyme levels are significantly lower in ribozyme ASO experiments, suggesting that the RT-PCR reaction is blocked by the ASO. ***B***, Ribozyme inhibition by ASO resulted in upregulation of *CPEB3* mRNA (exons 2–3). ***C***, Inhibition of CPEB3 ribozyme by ASO resulted in upregulation of *CPEB3* mRNA basal levels for exons 3–6 at the 2-hour time point. ***D***, Levels of exons 6–9 did not increase significantly at the 2-hour time point. ***E***, No statistically significant difference in CPEB3 ribozyme expression was observed after 24 hours post KCl induction, suggesting that all intronic RNA levels reached basal levels. ***F – H***, *CPEB3* mRNA expression largely returned to the basal level 24 hours post-stimulation, although levels of spliced exons 3–6 remained elevated (***F***: exons 2–3, ***G***: exons 3–6, ***H***: exons 6–9). ***I***, ASO treatment leads to an increase of *CPEB3* full-length mRNA (exons 2–9). ***J,*** qRT-PCR analysis of CPEB3 4^th^ intron expression reveals that the ribozyme ASO does not affect its levels, suggesting that it is specific for the ribozyme. ***K***, CPEB3 ribozyme ASO does not alter CPEB1, CPEB2, and CPEB4 mRNA expression, demonstrating the specificity of the ASO. ***L***, Effect of ASO treatment on cell viability. XTT assay was performed after 18 hours incubation of ASOs. Relative cell viability was normalized to the vehicle control.

To determine whether the ASO specifically targets CPEB3 ribozyme or modulates intron levels in general, we measured the levels of the 4^th^ CPEB3 intron, which does not harbor a self-cleaving ribozyme. No significant difference in the 4^th^ intron expression was observed between groups, demonstrating that the ASO does not have a broad non-specific effect on the stability of other introns (two-way ANOVA with Sidak’s *post hoc* tests, no significant effect of KCl: *F*_(1,18)_ = 4.187, *P* = 0.0566; no significant effect of ASO: *F*_(1,18)_ = 1.032, *P* = 0.3232; no significant interaction: *F*_(1,18)_ = 0.00001455, *P* = 0.9970; Fig. 3J). Similarly, we measured mRNA expression of other members of the *CPEB* gene family (*CPEB1*, *CPEB2*, and *CPEB4*), and our results revealed no significant difference in the gene expression between scrambled ASO and ASO groups (*CPEB1*: *t*_(8,777)_ = 0.6338, *P* = 0.5423; *CPEB2*: *t*_(7,768)_ = 1.491, *P* = 0.1753; *CPEB4*: *t*_(8.270)_ = 0.6268, *P* = 0.5477; unpaired *t* test; Fig. 3K). These results confirm that the ASO is specific for the CPEB3 ribozyme and only modulates levels of the *CPEB3* mRNA. To assess whether the ASO induces cytotoxicity *in vitro*, neuronal cultures were treated with either ASO or scrambled ASO. Cell viability was measured with an XTT assay and revealed no difference in either ASO-or scrambled-ASO-treated cells, compared to untreated cells. Thus, the ASOs used in this study did not induce cytotoxic effects in cultured neurons (scrambled ASO: *t*_(2.986)_ = 0.1257, *P* = 0.9079; ASO: *t*_(5.437)_ = 0.5869, *P* = 0.5808; unpaired *t* test; Fig. 3L).

### Ribozyme inhibition leads to increased expression of CPEB3 and plasticity-related proteins

We next determined whether inhibition of CPEB3 ribozyme affects CPEB3 protein expression. Treatment with the ribozyme ASO resulted in a significant increase in CPEB3 protein levels both in the basal state and under KCl-stimulated conditions, indicating a coordination of activity-dependent transcription and translation upon inhibition of CPEB3 ribozyme (two-way ANOVA with Sidak’s *post hoc* tests, significant effect of ASO: *F*_(1,24)_ = 21.68, *P* < 0.0001; no significant effect of KCl: *F*_(1,24)_ = 0.6204, *P* = 0.4386; no significant interaction: *F*_(1,24)_ = 1.556, *P* = 0.2243; Fig. 4A, B).

**Figure 4.**
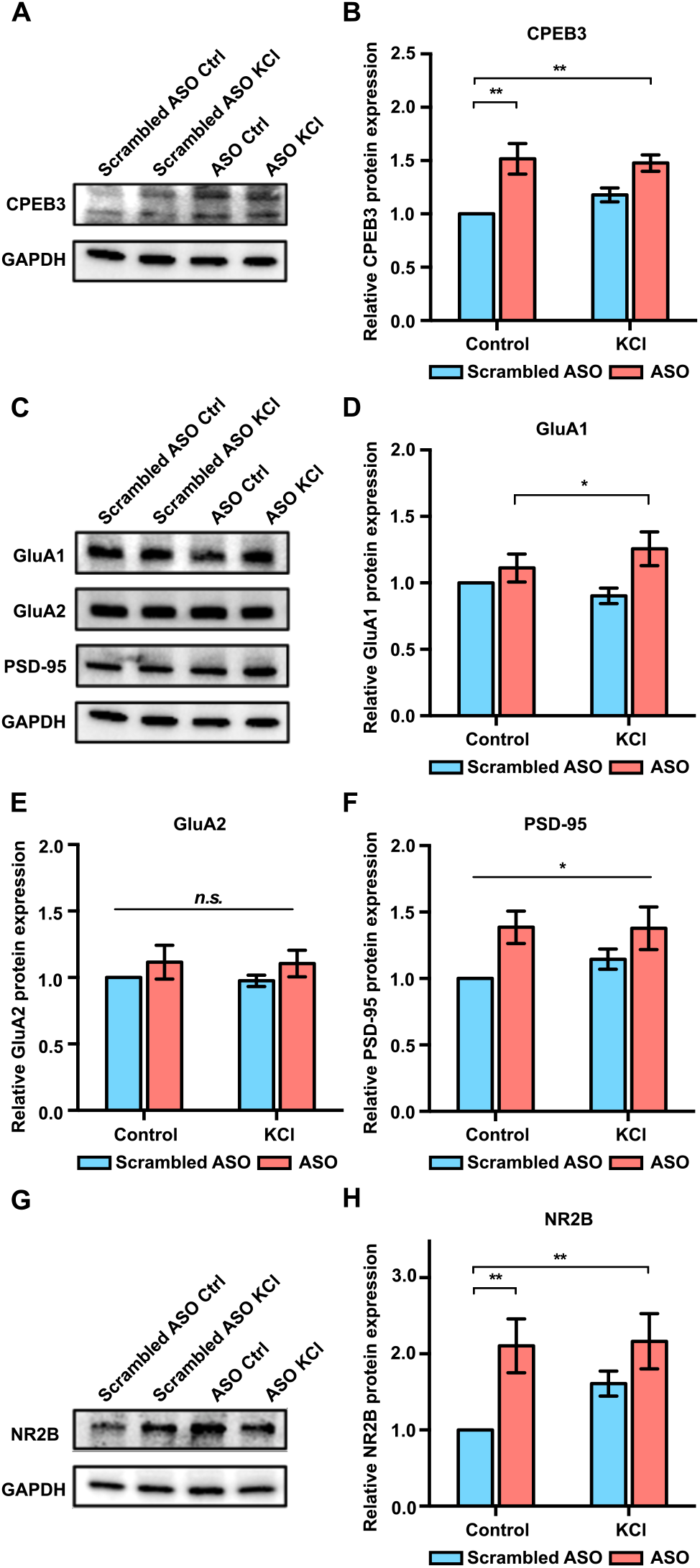
Effect of CPEB3 ribozyme ASO on protein expression in cultured cortical neurons (DIV7). ***A*,** Effect of CPEB3 ribozyme ASO on CPEB3 protein expression. Representative image of CPEB3 protein expression. GAPDH is used as a loading control. ***B***, Quantification of CPEB3 protein expression. Treatment of ASO followed by KCl stimulation led to an increase of CPEB3. ***C***, Representative immunoblotting image of GluA1, GluA2, and PSD-95 protein expression. GAPDH is used as a loading control. ***D***, Quantification of GluA1 protein expression. GluA1 is upregulated in the presence of ASO combined with neuronal stimulation. ***E***, Treatment with ASO leads to an increase of PSD-95 protein level in primary cortical neurons. ***F***, Quantification of GluA2 protein expression. No significant difference was observed between ASO and KCl groups. ***G***, Representative images of immunoblotting analysis showing NR2B protein expression. GAPDH is used as a loading control. ***H***, Quantification of NR2B protein expression. ASO treatment induces an increase in NR2B expression. \**P* < 0.05, \*\**P* < 0.01, *n.s.* not significant. Data are presented as mean ± SEM.

Previous studies have demonstrated the role of CPEB3 in the translational regulation of a number of plasticity-related proteins (PRPs), including AMPA-type glutamate receptors (AMPARs), NMDA receptor (NMDAR), and postsynaptic density protein 95 (PSD-95) (Huang et al., 2006; Chao et al., 2012; Chao et al., 2013; Fioriti et al., 2015). As an RNA-binding protein, CPEB3 binds to 3*’* UTR of GluA1, GluA2, and *PSD-95* mRNAs and regulates their polyadenylation and translation (Huang et al., 2006; Pavlopoulos et al., 2011; Chao et al., 2013; Fioriti et al., 2015). Treatment with the CPEB3 ribozyme ASO resulted in a significant increase in GluA1 and PSD-95 protein expression, whereas GluA2 levels remained unchanged (GluA1: two-way ANOVA with Sidak’s *post hoc* tests, significant effect of ASO: *F*_(1,24)_ = 7.134, *P* = 0.134; no significant effect of KCl: *F*_(1,24)_ = 0.07449, *P* = 0.7872; and no significant interaction: *F*_(1,24)_ = 1.911, *P* = 0.1796; Fig. 4C and 4D. GluA2: two-way ANOVA with Sidak’s *post hoc* tests, no significant effect of ASO: *F*_(1,24)_ = 2.149, *P* = 0.1556; no significant effect of KCl: *F*_(1,24)_ = 0.04578, *P* = 0.8324; and no significant interaction: *F*_(1,24)_ = 0.006228, *P* = 0.9358; Fig. 4C and 4E. PSD-95: two-way ANOVA with Sidak’s *post hoc* tests, significant effect of ASO: *F*_(1,24)_ = 8.213, *P* = 0.0085; no significant effect of KCl: *F*_(1,24)_ = 0.4082, *P* = 0.5290; and no significant interaction: *F*_(1,24)_ = 0.5106, *P* = 0.4818; Fig. 4C and 4F). Likewise, ASO treatment led to an upregulation of NR2B protein, which is one of the NMDAR subunits (two-way ANOVA with Sidak’s *post hoc* tests, significant effect of ASO: *F*_(1,19)_ = 10.40, *P* = 0.0045; no significant effect of KCl: *F*_(1,19)_ = 1.791, *P* = 0.2078; and no significant interaction: *F*_(1,19)_ = 1.444, *P* = 0.2982; Fig. 4G and 4H). Thus, our results demonstrate that CPEB3 ribozyme activity affects several downstream processes, particularly mRNA maturation and translation, but also the expression of PRPs, including the translation of AMPAR and NMDAR mRNAs.

### CPEB3 ribozyme ASO leads to an increase of *CPEB3* mRNA and polyadenylation of PRPs in the CA1 hippocampus

To investigate whether the CPEB3 ribozyme exhibits similar effects in regulating genes related to synaptic plasticity *in vivo*, mice were stereotaxically infused with either ribozyme ASO, scrambled ASO, or vehicle into the CA1 region of the dorsal hippocampus, a major brain region involved in memory consolidation and persistence (Fig. 5A). Infusion of the ASO targeting the CPEB3 ribozyme significantly reduced ribozyme levels detected by RT-qPCR in the dorsal hippocampus (one-way ANOVA with Sidak’s *post hoc* tests; *F*_(2,18)_ = 3.901, *P* = 0.0391; Fig. 5B). However, administration of ASO led to an increase of *CPEB3* mRNA in the CA1 hippocampus (one-way ANOVA with Sidak’s *post hoc* tests; exons 2–3: *F*_(2,18)_ = 6.199, *P* = 0.0089; exons 3–6: *F*_(2,18)_ = 12.44, *P* = 0.0004; exons 6–9: *F*_(2,17)_ = 11.03, *P* = 0.0008; Fig. 5C), confirming that the ASO prevents ribozyme self-scission during CPEB3 pre-mRNA transcription and thereby increases *CPEB3* mRNA levels. To further determine the effect of CPEB3 ribozyme in regulating mature mRNA processing, the level of *CPEB3* exons 2–9 was measured. ASO-infused mice exhibited a significant increase in full-length *CPEB3* mRNA (one-way ANOVA with Sidak’s *post hoc* tests; *F*_(2,17)_ = 4.385, *P* = 0.0291; Fig. 5D). In line with our *in vitro* studies, no significant difference in the ribozyme-free 4^th^ intron levels was observed between mouse hippocampus treated with ASO and vehicle (one-way ANOVA with Sidak’s *post hoc* tests; *F*_(2,18)_ = 0.3663, *P* = 0.6984; Fig. 5E). We also found no significant difference in levels of other *CPEB* mRNAs or degree of protein expression between ASO and control groups (one-way ANOVA with Sidak’s *post hoc* tests; *CPEB1* mRNA: *F*_(2,18)_ = 0.8203, *P* = 0.4570; Fig. 5F; *CPEB2* mRNA: *F*_(2,18)_ = 2.002, *P* = 0.1641; Fig 5F; *CPEB4* mRNA: *F*_(2,18)_ = 0.3562, *P* = 0.7052; Fig 5F; CPEB1 protein: *t*_(8.942)_ = 0.4469, *P* = 0.6656; Fig. 5G and 5H. CPEB4 protein: *t*_(10.24)_ = 1.089, *P* = 0.3012; Fig. 6G and 6H). These findings demonstrate that the ASO used in this study targets the CPEB3 ribozyme *in vivo* with high specificity.

**Figure 5.**
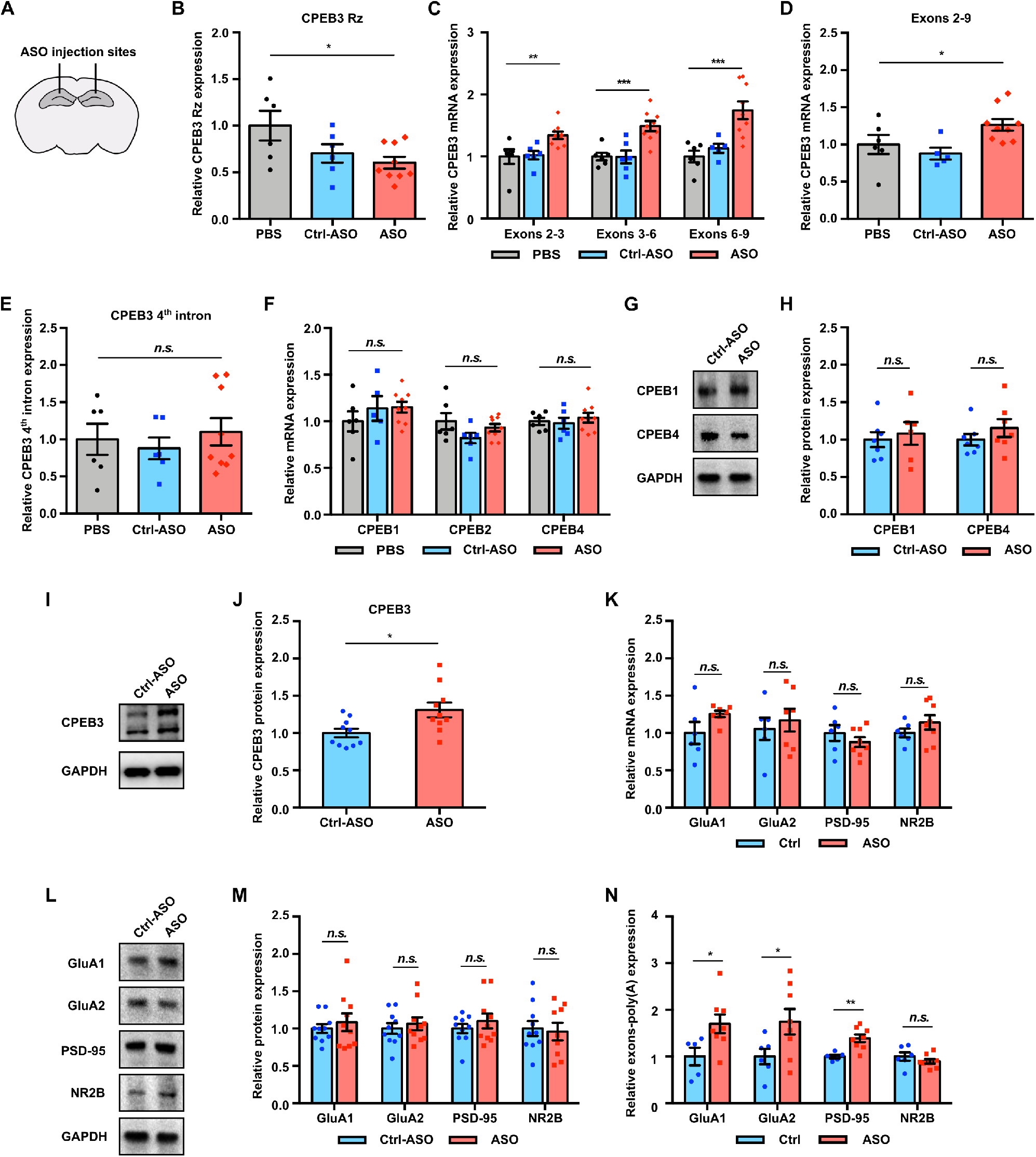
CPEB3 ribozyme ASO leads to an increase of *CPEB3* mRNA and polyadenylation of PRPs in the CA1 hippocampus. ***A***, Schematic representation of stereotaxic procedure. ASO, scrambled ASO, or vehicle was bilaterally infused to the mouse CA1 hippocampus. ***B***, Validation of CPEB3 ribozyme knockdown *in vivo*. Administration of CPEB3 ribozyme ASO to the mouse CA1 hippocampus leads to a decrease in CPEB3 ribozyme levels. ***C***, *CPEB3* mRNA expression is upregulated in the CPEB3 ribozyme ASO treatment group compared to controls. ***D***, *CPEB3* full-length mRNA (exons 2–9) is significantly elevated in the presence of ASO. ***E***, The CPEB3 ribozyme ASO has high specificity for its cleavage site (in the 3^rd^ intron) *in vivo*. qRT-PCR analysis of the 4^th^ intron of *CPEB3* gene demonstrates no significant difference between controls and ASO groups. ***F***, qRT-PCR analysis reveals no significant difference between controls and ASO groups in *CPEB1*, *CPEB2*, and *CPEB4* mRNA expression. ***G***, Effect of CPEB3 ribozyme on CPEB1 and CPEB4 protein expression. GAPDH is used as a loading control. ***H***, Quantification of CPEB1 and CPEB4 protein expression. CPEB3 ribozyme ASO does not change CPEB1 and CPEB4 protein expression. ***I***, Effect of CPEB3 ribozyme on CPEB3 protein expression. Representative image of immunoblotting analysis. GAPDH is used as a loading control. ***J***, Quantification of CPEB3 protein expression. CPEB3 ribozyme ASO leads to an increase of CPEB3 protein expression in the CA1 hippocampus. ***K***, Inhibition of CPEB3 ribozyme does not affect transcription of other plasticity-related genes. qRT-PCR analysis of mature GluA1, GluA2, PSD-95, and NR2B mRNAs. No significant difference between ASO and control was observed for splice junctions within the mRNAs, showing that modulation of the CPEB3 ribozyme does not affect transcription or splicing of these mRNAs. ***L***, Effect of CPEB3 ribozyme on PRP protein expression. Representative images of immunoblotting analysis. GAPDH is used as a loading control. ***M***, Quantification of PRP protein expression. Blocking CPEB3 ribozyme does not affect PCPs protein expression in the naïve state. ***N***, Inhibition of CPEB3 ribozyme resulted in increased polyadenylation of plasticity-related genes. \**P* < 0.05, \*\**P* < 0.01, \*\*\**P* < 0.001, *n.s.* not significant. Data are presented as mean ± SEM.

**Figure 6.**
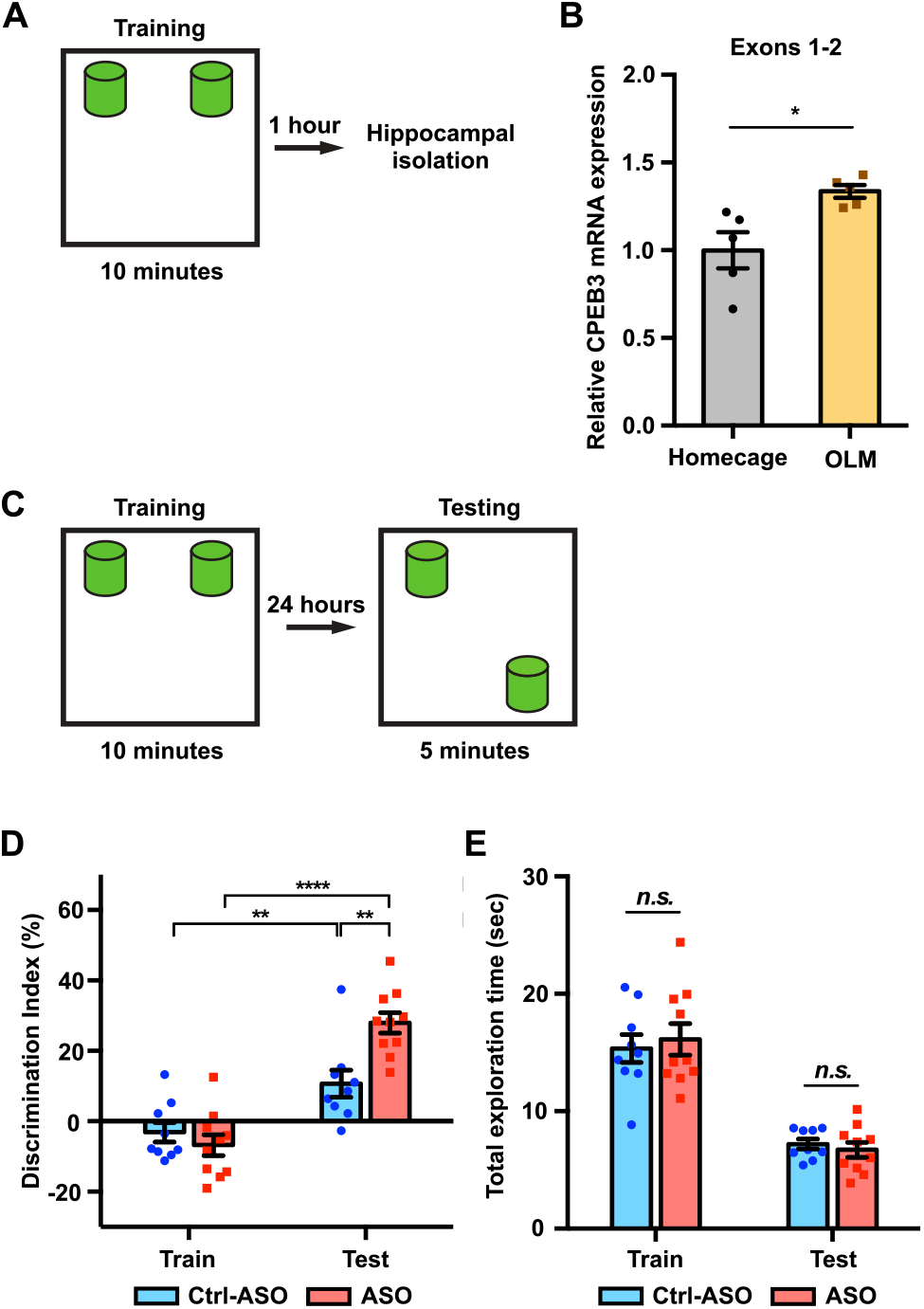
Inhibition of CPEB3 ribozyme enhances long-term OLM. ***A***, Schematic representation of how the hippocampal gene expression is examined after OLM training task. ***B***, OLM training induces upregulation of *CPEB3* mRNA in the CA1 hippocampus. ***C***, Experimental procedure testing long-term memory. ***D***, Mice infused with scrambled ASO or CPEB3 ribozyme ASO showed no preference for either object in OLM training. Mice infused with CPEB3 ribozyme ASO show significant discrimination index in OLM testing. ***E***, CPEB3 ribozyme ASO and control mice display similar total exploration time. \**P* < 0.05, \*\**P* < 0.01, \*\*\*\**P* < 0.0001, *n.s.* not significant. Data are presented as mean ± SEM.

Next, we tested whether the CPEB3 ribozyme inhibition affects *CPEB3* translation. The elevated CPEB3 protein expression found in ASO-treated mice suggests that increased translation of *CPEB3* directly results from increased levels of full-length mRNA (*t*_(14.50.)_ = 2.709, *P* = 0.0165; unpaired *t* test; Fig. 5I, 5J). Furthermore, blocking the CPEB3 ribozyme does not change GluA1, GluA2, PSD-95, and NR2B mRNA or protein expression in naïve, home cage mice (GluA1: *t*_(5.848)_ = 1.655, *P* = 0.1503; GluA2: *t*_(10.96)_ = 0.5476, *P* = 0.5949; PSD-95: *t*_(8.760)_ = 0.9838, *P* = 0.3516; NR2B: *t*_(11.11)_ = 1.250, *P* = 0.2369; Fig. 5K; GluA1: *t*_(13.18)_ = 0.6339, *P* = 0.5370; GluA2: *t*_(17.54)_ = 0.5755, *P* = 0.5723; PSD-95: *t*_(14.94)_ = 0.8612, *P* = 0.4027; NR2B: *t*_(16.34)_ = 0.2604, *P* = 0.7978; unpaired *t* test; Fig. 5L, 5M). Thus, in naïve mice, ribozyme inhibition leads to increased basal levels of the *CPEB3* mRNA and protein, but its downstream mRNA targets remain unchanged in the absence of activity-dependent learning or stimulation.

The CPEB3 ribozyme activity may result from polyadenylation of its target mRNAs, and therefore, 3′ rapid amplification of cDNA ends (3′ RACE) was performed to examine the 3′ termini of several mRNAs. We found that ribozyme ASO administration led to increased GluA1, GluA2, and PSD-95 mRNA polyadenylation in the mouse dorsal hippocampus (GluA1: *t*_(10.44)_ = 2.535, *P* = 0.0287; GluA2: *t*_(11.02)_ = 2.327, *P* = 0.0400; PSD-95: *t*_(9.808)_ = 4.254, *P* = 0.0018; NR2B: *t*_(8.020)_ = 0.9846, *P* = 0.3536; unpaired *t* test; Fig. 5N). These data support a model wherein the inhibition of the CPEB3 ribozyme leads to increased polyadenylation of existing AMPARs and PSD-95 mRNAs, and suggests a role for the ribozyme in post-transcriptional regulation and 3′ mRNA processing.

### Inhibition of CPEB3 ribozyme in the dorsal hippocampus enhances long-term memory

Previous studies have shown that CPEB3 is regulated by synaptic activity; for example, Morris water maze (MWM) training and contextual fear conditioning induced an increase in CPEB3 protein expression, and *CPEB3* mRNA was upregulated 2 hours after kainate injection (Theis et al., 2003). To examine whether *CPEB3* mRNA is modulated by behavioral training, we subjected mice to an object location memory (OLM) task (Vogel-Ciernia and Wood, 2014) and isolated hippocampal tissues 1 hour after training (Fig. 6A). The OLM task has been widely used to study hippocampal-dependent spatial memory. The task is based on an animal’s innate preference for novelty and its capability for discriminating spatial relationships between novel and familiar object locations (Vogel-Ciernia and Wood, 2014). We first examined the effect of training on *CPEB3* mRNA expression. *CPEB3* mRNA exons 1–2, which span about 33 kb of the gene downstream of the promoter (Fig. 1A), were upregulated 1 hour after training compared to naïve mice (Exons 2–3: *t*_(4.991)_ = 3.085, *P* = 0.0274; Fig. 6B). Although a previous study reported that the *CPEB3* mRNA level (exons 2– 6) was not altered after a MWM test, these seemingly contradictory results can be explained by the time points and segments of the mRNA analyzed. The distance from the 5ʹ terminus of the pre-mRNA and exon 2 is about 33 kb, whereas exon 6 is more than three times farther (110 kb). As a result, RNAP II and the splicing machinery require at least three times longer to produce the spliced exons 2–6 of the *CPEB3* mRNA (assuming no significant pausing in transcription and co-transcriptional splicing). Transcription initiation, pre-mRNA production up to exon 2, and splicing would be expected to yield spliced mRNA exons 1–2 after 1 hour, but reaching the 6^th^ exon and splicing the mRNA would likely not happen in that time frame. We therefore believe the results of these two studies are not at odds; rather, these results demonstrate that the detection of new rounds of gene expression should rely on measurements of early segments of activity-induced genes, rather than later segments.

To assess whether inhibition of the CPEB3 ribozyme improves memory formation, we studied the effect of the ASO on long-term memory formation for object location using the OLM task (Fig. 6C). We infused mice bilaterally into the CA1 dorsal hippocampus with the CPEB3 ribozyme ASO, scrambled ASO, or vehicle 48 hours prior to OLM training. Mice exhibit no preference for either object, as demonstrated the absence of significant difference in training discrimination index (DI) (*t*_(16.99)_ = 0.8967, *P* = 0.3824; unpaired *t* test; Fig. 6D). Likewise, during training and testing sessions, similar total exploration times were observed for ASO-infused mice and control mice, demonstrating that both groups of mice have similar exploitative behavior and that the ASO did not simply affect locomotor or exploration performance (Train: *t*_(17.00)_ = 0.2342, *P* = 0.8176, Test: *t*_(13.48)_ = 1.644, *P* = 0.1232; unpaired *t* test; Fig. 6E). However, the CPEB3 ribozyme ASO mice showed a significant increase in DI between training and testing compared to control groups, suggesting that these mice experienced a robust enhancement of novel object exploration (ASO × session interaction *F*_(1,34)_ = 11.06, *P* = 0.0021; two-way ANOVA with Sidak’s *post hoc* tests; Fig. 6D). These results provide strong evidence that *CPEB3* is critical for long-term memory, and that the CPEB3 ribozyme activity is anti-correlated with the formation of long-term memory.

### CPEB3 ribozyme ASO leads to an upregulation in protein expression of CPEB3 and PRPs during memory consolidation

Learning-induced changes in gene expression and protein synthesis are essential for memory formation and consolidation (Kandel, 2001). To determine whether upregulation of *CPEB3* mRNA by the ribozyme ASO leads to a change in expression of the CPEB3 protein and its downstream targets, we analyzed the dorsal hippocampal homogenates and synaptosomal fractions. Administration of CPEB3 ribozyme ASO led to a significant increase of CPEB3 protein expression in the CA1 hippocampal homogenates and crude synaptosomes 1 hour after OLM testing (hippocampal homogenates: *t*_(17.00)_ = 2.345, *P* = 0.0314; crude synaptosomes: *t*_(11.11)_ = 2.403, *P* = 0.0349; unpaired *t* test; Fig 7A, B, D). This result confirms that ASO-mediated knockdown of the CPEB3 ribozyme facilitates *CPEB3* mRNA processing and translation. In addition, the protein levels of GluA1, GluA2, PSD-95, and NR2B were measured to determine whether increased CPEB3 further regulates translation of PRPs. In total tissue lysates, no significant difference in PRP levels was observed between ASO and control (GluA1: *t*_(15.96)_ = 0.3751, *P* = 0.7125; GluA2: *t*_(15.16)_ = 0.9432, *P* = 0.3604; PSD-95: *t*_(17.63)_ = 0.2849, *P* = 0.7790; NR2B: *t*_(17.32)_ = 0.9415, *P* = 0.3594; unpaired *t* test; Fig. 7A, C). However, in synaptosomal fractions, GluA1, PSD-95, and NR2B protein levels were increased in ASO-infused mice, relative to scrambled ASO control animals; the GluA2 protein level was unaffected (GluA1: *t*_(15.83)_ = 2.433, *P* = 0.0272; GluA2: *t*_(14.40)_ = 1.497, *P* = 0.1559; PSD-95: *t*_(17.25)_ = 2.115, *P* = 0.0493; NR2B: *t*_(12.42)_ = 3.174, *P* = 0.0077; unpaired *t* test; Fig. 7, A, E). Our findings thus show that blocking CPEB3 ribozyme activity leads to an increase in CPEB3 protein production, and upregulation of CPEB3 by OLM further causes an increase in local GluA1, PSD-95, and NR2B translation.

**Figure 7.**
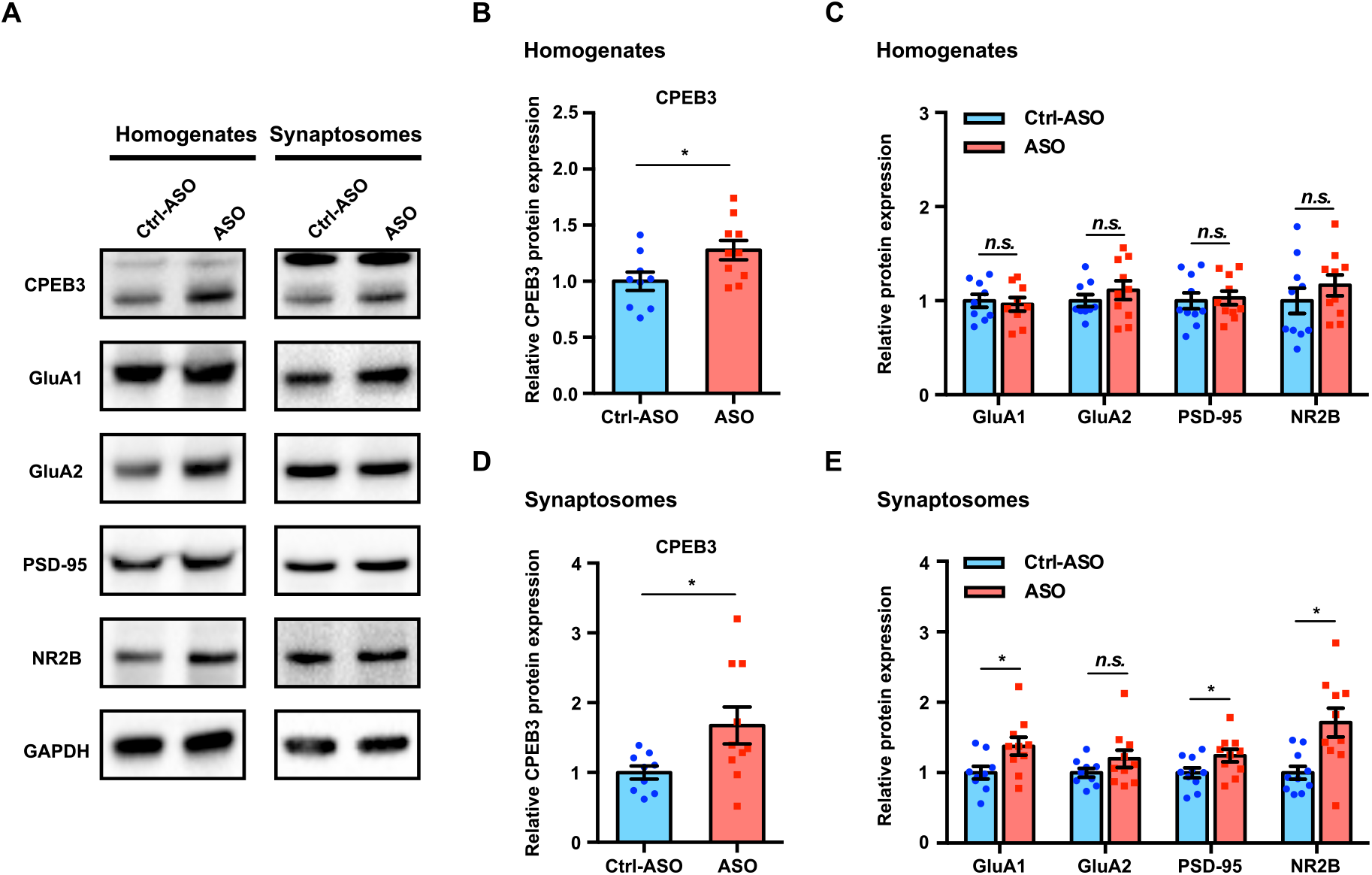
Inhibition of CPEB3 ribozyme leads to upregulation of CPEB3 and PRPs protein expression after OLM. ***A***, Representative images of immunoblotting analysis. GAPDH is used as a loading control. ***B – C***, Quantification of CPEB3 (**B**) and PRPs (**C**) in tissue homogenates shows increased expression of CPEB3 but not of PRPs. ***D – E***, In synaptosomes, the protein expression of both CPEB3 (**D**) and PRPs (**E**) is increased. \**P* < 0.05, *n.s.* not significant. Data are presented as mean ± SEM.

## Discussion

Self-cleaving ribozymes are broadly distributed small functional RNAs that promote an intramolecular, site-specific, self-scission reaction (Buzayan et al., 1986; Hutchins et al., 1986; Prody et al., 1986; Sharmeen et al., 1988; Saville and Collins, 1990; Jimenez et al., 2015). Despite distinct structures and cut sites, these natural self-cleaving ribozymes all accelerate the same transesterification reaction, which operates via an acid–base catalysis mechanism: nucleophilic attack of a ribose 2′-oxyanion on the adjacent phosphodiester bond yields a 2′,3′-cyclic phosphate and a 5′-hydroxyl product (Wu et al., 1989; Fedor, 2009; Jimenez et al., 2015; Wilson et al., 2016; Ren et al., 2017; Seith et al., 2018; Peng et al., 2021). Self-cleaving ribozymes act in *cis* (i.e., cut their own backbone) and therefore execute a single catalytic turnover. To date, 10 distinct families of self-cleaving ribozymes have been discovered (Peng et al., 2021), but relatively little is known about their biological roles.

The HDV family of ribozymes has been extensively studied: crystal structures have been elucidated, and the mechanism of self-scission (based on a general acid–base catalysis) is well-established (Ferre-D’Amare et al., 1998; Ke et al., 2004; Das and Piccirilli, 2005; Chen et al., 2010; Koo et al., 2015). These ribozymes operate during rolling circle replication of the HDV RNA genome and in processing of certain non-LTR retrotransposons (Sharmeen et al., 1988; Wu et al., 1989; Eickbush and Eickbush, 2010; Ruminski et al., 2011; Sanchez-Luque et al., 2011), but given their broad distribution in nature, their biological roles remain largely unexplored. Mammals harbor several self-cleaving ribozymes, all with unknown biological functions (Salehi-Ashtiani et al., 2006; Martick et al., 2008; de la Pena and Garcia-Robles, 2010; Perreault et al., 2011; Hernandez et al., 2020; Chen et al., 2021). One of these ribozymes, the HDV-like CPEB3 ribozyme, which is a functionally conserved self-cleaving RNA (Bendixsen et al., 2021), maps to the second intron of the *CPEB3* gene (Fig. 1A), and its *in vitro* activity (Fig. 1B) suggests that its self-scission may be tuned to disrupt the intron at a rate that is similar to the production speed of the downstream intronic sequence ahead of the next exon. Given that the self-scission of intronic ribozymes is inversely correlated with splicing efficiency of the harboring pre-mRNA (Fong et al., 2009), we investigated how the endogenous intronic ribozyme affects the *CPEB3* mRNA maturation and translation, and how it affects memory formation in mice.

Modifications of synaptic strength are thought to underlie learning and memory in the brain. Studies in hippocampal slices revealed local translation in dendrites following induction of LTP (Frey and Morris, 1997). Cytoplasmic polyadenylation-induced translation is one of the key steps critical to controlling protein synthesis and neuroplasticity (Du and Richter, 2005; Richter, 2007, 2010), and one of the proteins involved in regulating cytoplasmic polyadenylation of mRNAs is CPEB3. In *Aplysia* sensory-motor neuron co-culture, application of repeated pulses of serotonin (5-HT) induces ApCPEB protein expression at the stimulated synapses and, as a result, LTF, which is a form of learning-related synaptic plasticity that is widely studied in *Aplysia* (Si et al., 2003; Si et al., 2010). In murine primary hippocampal neurons, the level of CPEB3 protein expression is positively regulated by neuronal activity (Fioriti et al., 2015) and plays dual roles in regulating mRNA translation (Du and Richter, 2005; Stephan et al., 2015): a post-translational modification of CPEB3 (monoubiquitination by Neuralized1) converts it from a repressor to an activator (Pavlopoulos et al., 2011).

Several studies have shown that CPEB3 is essential for synaptic strength, regulating mRNA translation of several PRPs at synapses (Huang et al., 2006; Pavlopoulos et al., 2011; Fioriti et al., 2015). Previous reports have shown that CPEB3 regulates GluA1 and GluA2 polyadenylation: *CPEB3* conditional knockout mice fail to elongate the poly(A) tail of GluA1 and GluA2 mRNA after Morris water maze training, and overexpression of CPEB3 changes the length of the GluA1 and GluA2 mRNA poly(A) tail (Fioriti et al., 2015). Hippocampal-dependent learning and memory is modulated by CPEB3 on the level of translation (Pavlopoulos et al., 2011), but it is unknown whether the CPEB3 expression is modulated by the CPEB3 ribozyme.

In mammals, the coordination of pre-mRNA processing and transcription can affect gene expression (Neugebauer, 2019). Using long-read sequencing and Precision Run-On sequencing (PRO-seq) approaches, measurements of co-transcriptional splicing events in mammalian cells demonstrated that co-transcriptional splicing efficiency impacts productive gene output (Reimer et al., 2021). The temporal and spatial window shows that the splicing and transcription machinery are tightly coupled. Our study is agreement with this co-transcriptional splicing model and shows that inhibition of the intronic CPEB3 ribozyme leads both to an increase in *CPEB3* mRNA and protein levels in primary cortical neurons and the dorsal hippocampus upon synaptic stimulation, and subsequently, to changes in the polyadenylation of target mRNAs of the CPEB3 protein.

Activity-dependent synaptic changes are governed by AMPAR trafficking, and AMPARs are mobilized to the post-synaptic surface membrane in response to neuronal activity in a dynamic process (Diering and Huganir, 2018). Our data demonstrate that the activation of CPEB3 by neuronal stimulation further facilitates translation of PRPs *in vivo*. These observations are consistent with a model in which learning induces CPEB3 protein expression, and ablation of CPEB3 abolishes the activity-dependent translation of GluA1 and GluA2 in the mouse hippocampus (Fioriti et al., 2015). Specifically, it has been suggested that CPEB3 converts to prion-like aggregates in stimulated synapses that mediate hippocampal synaptic plasticity and facilitate memory storage (Si and Kandel, 2016). Because training can produce effective long-term memory, it is likely that increased CPEB3 protein expression due to CPEB3 ribozyme inhibition further facilitates experience-induced local translational processes.

ASOs have been used in many studies to inhibit specific mRNAs. A notable example is an FDA-approved ASO that modulates co-transcriptional splicing of the *SMN2* mRNA (Hua et al., 2010). More recently, Tran *et al*. demonstrated that ASO can suppress hexonucleotide repeat expansion of the first intron in the *C9ORF72* gene (Tran et al., 2022). Our work shows that an ASO designed to bind the substrate strand of an endogenous self-cleaving ribozyme (located in an intron) increases the expression of the fully spliced mRNA that harbors the ribozyme. Interestingly, our experiments with inhibitory ASO yielded lower ribozyme levels than control experiments, suggesting that the ASO directs degradation of the target sequence; however, this degradation must occur on a timescale that is longer than the splicing of the mRNA, because we consistently measure higher mRNA levels when the ribozyme is inhibited. Given that three endogenous mammalian self-cleaving ribozymes map to introns (Salehi-Ashtiani et al., 2006; de la Pena and Garcia-Robles, 2010; Perreault et al., 2011), we anticipate that application of our ASO strategy will help to decipher the effect of these self-cleaving ribozymes on their harboring mRNAs and to elucidate their biological roles.

In summary, our study describes a unique role for the CPEB3 ribozyme in post-transcriptional maturation of *CPEB3* mRNA and its subsequent translation in mouse CA1 hippocampus. Inhibition of the CPEB3 ribozyme by ASO and OLM training induce activity-dependent upregulation of CPEB3 and local production of PRPs. These molecular changes are critical for establishing persistent changes in synaptic plasticity that are required for long-term memory. Thus, our study has identified a novel biological role for self-cleaving ribozymes in the brain. More broadly, we have demonstrated a method for determining the biological roles of self-cleaving ribozymes in both mammals (as shown here) and other organisms.

## Acknowledgments

We thank M. Malgowska, C-K. Lau, and M. A. Sta Maria for experimental assistance, and L. Fioriti and E. Kandel for encouragement and support during early parts of the project.

## Author contributions

Design of cell culture experiments: CCC, XL, TWB, AL

Design of mouse experiments: CCC, MAW, AL

In vitro ribozyme kinetics measurements: MM

Design of ASOs: MN

Cell culture experiments: CCC, LT, XL

Mouse experiments: CCC

Stereotaxic surgeries and in vivo behavior experiments: JH, CC

Data analysis: CCC

Writing—original draft: CCC, AL

Writing—review & editing: CCC, MN, XL, LT, TWB, MAW, AL

